# The one-message-per-cell-cycle rule: A conserved minimum transcription level for essential genes

**DOI:** 10.1101/2023.07.06.548020

**Authors:** Teresa W. Lo, Han Kyou James Choi, Dean Huang, Paul A. Wiggins

**Affiliations:** Department of Physics, University of Washington, Seattle, Washington 98195, USA; Department of Bioengineering, University of Washington, Seattle, Washington 98195, USA; Department of Microbiology, University of Washington, Seattle, Washington 98195, USA

## Abstract

The inherent stochasticity of cellular processes leads to significant cell-to-cell variation in protein abundance. Although this noise has already been characterized and modeled, its broader implications and significance remain unclear. In this paper, we revisit the noise model and identify the number of messages transcribed per cell cycle as the critical determinant of noise. In yeast, we demonstrate that this quantity predicts the non-canonical scaling of noise with protein abundance, as well as quantitatively predicting its magnitude. We then hypothesize that growth robustness requires an upper ceiling on noise for the expression of essential genes, corresponding to a lower floor on the transcription level. We show that just such a floor exists: a minimum transcription level of one message per cell cycle is conserved between three model organisms: *Escherichia coli*, yeast, and human. Furthermore, all three organisms transcribe the same number of messages per gene, per cell cycle. This common transcriptional program reveals that robustness to noise plays a central role in determining the expression level of a large fraction of essential genes, and that this fundamental optimal strategy is conserved from *E. coli* to human cells.

## INTRODUCTION

All molecular processes are inherently stochastic on a cellular scale, including the processes of the central dogma, responsible for gene expression [1, 2]. As a result, the expression of every protein is subject to cell-to-cell variation in abundance [1]. Many interesting proposals have been made to describe the potential biological significance of this noise, including bet-hedging strategies, the necessity of feedback in gene regulatory networks, *etc* [1, 3, 4]. However, it is less clear to what extent noise plays a central role in determining the function of the gene expression process more generally. For instance, Hausser *et al*. have described how the tradeoff between economy (*e*.*g*. minimizing the number of transcripts) and precision (minimizing the noise) explains why genes with high transcription rates and low translation rates are not observed [5]. Although these results suggest that noise may provide some coarse limits on the function of gene expression, this previous work does not directly address a central challenge posed by noise: How does the cell ensure that the lowest expression essential genes, which are subject to the greatest noise, have sufficient abundance in all cells for robust growth?

To investigate this question, we first focus on noise in *Saccharomyces cerevisiae* (yeast), and find that the noise scaling with protein abundance is not canonical. We re-analyze the canonical stochastic kinetic model for gene expression, the telegraph model [6–8], to understand the relationship between the underlying kinetic parameters and the distribution of protein abundance in the cell. As previously reported, we find that the protein abundance for a gene is described by a gamma distribution with two parameters: the *message number*, defined as the total gene message number transcribed per cell cycle, and the translation efficiency, which is the mean protein number translated per message. Protein expression noise is completely determined by the message number [3, 9]. Although these results have been previously reported, the distinction between message number *per cell* versus *per cell cycle* and even between *mean protein number* and *mean message number* is often neglected (*e*.*g*. [10]).

To explore the distinction between these parameters and provide clear evidence of the importance of the message number, we return to the analysis of noise in yeast. In yeast, the translation efficiency increases with message number [11]. By fitting an empirical model for the translation efficiency, we demonstrate that the noise should scale with a half-power of protein abundance. We demonstrate that this non-canonical scaling is observed and that our translation model makes a parameter-free prediction for the noise. The prediction is in close quantitative agreement with observation [12], confirming that the message number is the key determinant of noise strength.

Finally, we use this result to explore the hypothesis that there is a minimum expression level for essential genes, dictated by noise. The same mean expression level can be achieved by a wide range of different translation and transcription rates with different noise levels. We hypothesize that growth robustness requires that essential genes (but not non-essential genes) are subject to a floor expression level, below which there is too much cell-to-cell variation to ensure growth. To test this prediction, we analyze transcription in three model organisms, *Escherichia coli*, yeast, and *Homo sapiens* (human), with respect to three related gene characteristics: transcription rate, cellular message number, and message number per cell cycle. As predicted by the noise-based mechanism, we observe an organism-independent floor for the number of messages transcribed per cell cycle for essential genes, but not non-essential genes. We conclude that virtually all essential genes are transcribed at a rate of at least once per cell cycle. This analysis strongly supports the hypothesis that the same biological optimization imperatives, which determine the transcription rates of many low-expression genes, are conserved from *E. coli* to human.

## RESULTS

### Implications of noise on growth robustness

With the realization of the stochasticity of central dogma processes, a key question is how cells can grow robustly in spite of cell-to-cell variations in protein expression. The noise in protein abundance is defined as the coefficient of variation squared [12–14]:

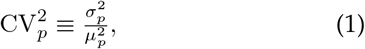

where 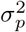 is the variance of protein number and 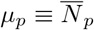 its mean. It is important to emphasize that protein abundance must double between birth and cell division in symmetrically dividing cells during steady state growth. The protein abundance should therefore be interpreted either as expression per unit volume [15] or the abundance associated with cells of a defined volume [12].

The coefficient of variation is inversely related to protein abundance and therefore low-copy proteins have the highest noise [3, 9, 12–15]. The challenge faced by the cell is that many essential proteins, strictly required for cell growth, are relatively low abundance. How does the cell ensure sufficient protein abundance in spite of cell-to-cell variation in protein number? It would seem that growth robustness demands that, for essential proteins, the mean should be greater than the standard deviation:

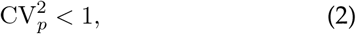

in order to ensure that protein abundance is sufficiently high enough to avoid growth arrest. To what extent do essential proteins obey this noise threshold?

### What determines the strength of the noise?

Usually, noise is argued to be proportional to inverse protein abundance (*e*.*g*. [3, 4, 10]):

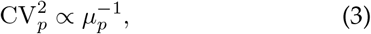

for low abundance proteins, motivated both by theoretical and experimental results [10, 15] and in some cases obeying a low-translation efficiency limit [15]:

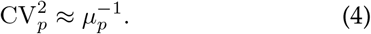

Can this model be used to make quantitative predictions of the noise? *E*.*g*., is the scaling of Eq. 3 correct? Can the coefficient of proportionality be predicted? Although Eq. 3 appears to describe *E. coli* quite well [15], the situation in yeast is more complicated [16]. To analyze the statistical significance of the deviation from the canonical noise model in yeast, we can fit an empirical model to the noise [13, 14]:

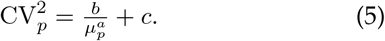

In the null hypothesis, *a* = 1 (canonical scaling), while *b* and *c* are unknown parameters. *c* corresponds to the noise floor. In the alternative hypothesis, all three coefficients are unknown. (A detailed description of the statistical model is given in the Supplemental Material Sec. A 5.)

The canonical model fails to fit the noise data for yeast as reported by Newman *et al*. [12]: The null hypothesis is rejected with p-value *p* = 6 × 10^*-*36^. The model fit to the data is shown in Fig. 2. The estimated scaling exponent for protein abundance in the alternative hypothesis is *a* = 0.57 *±* 0.02, and a detailed description of the statistical model and parameter fits is provided in Supplementary Material Sec. A5 d. As shown in Fig. 2, even from a qualitative perspective, the scaling of the yeast noise at low copy number is much closer to 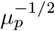 than to canonical assumption 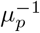 (Eq. 3). In particular, above the detection threshold, the noise is always larger than the low-translation efficiency limit (Eq. 4).

**FIG. 1.**
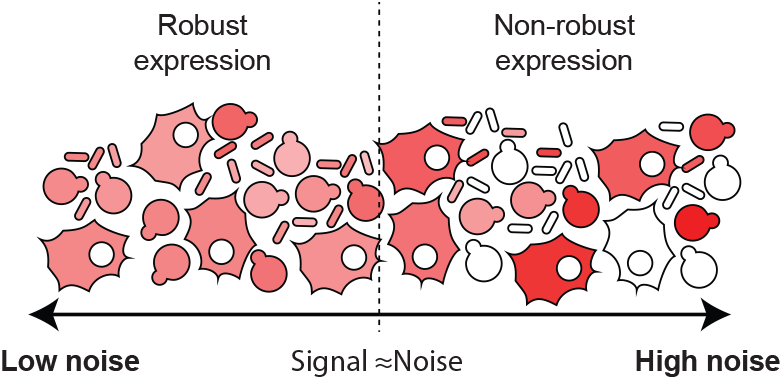
Robustness hypothesis: The stochasticity in gene expression is represented by the red shading. We hypothesize that robust growth requires sufficiently low noise levels for cellular function. We hypothesize that this critical noise level should be below the level where the signal (mean) equals noise (standard deviation).

**FIG. 2.**
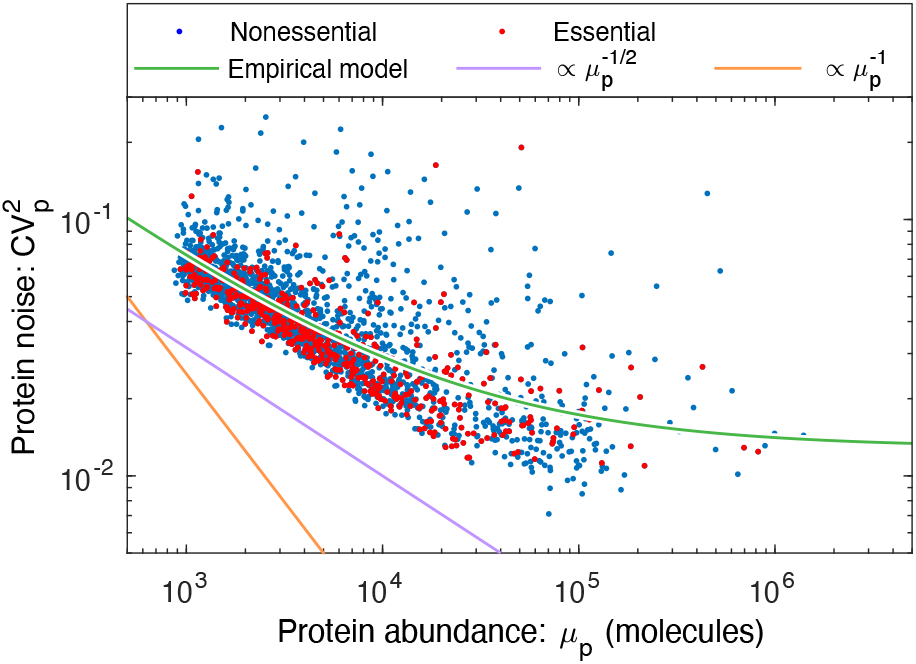
A non-canonical scaling is observed for gene-expression noise in yeast. The protein expression noise 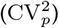 for yeast scales like 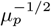 (purple) rather than the canonical 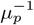 (orange) for low-abundance proteins. (Data from Ref. [12].) An empirical noise model (Eq. 5, green) fit to the essential genes gives an estimate of the protein-abundance scaling of 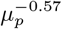.

### Stochastic kinetic model for central dogma

To understand the failure of the canonical assumptions, we revisit the underlying model. The telegraph model for the central dogma describes multiple steps in the gene expression process: Transcription generates mRNA messages [17]. These messages are then translated to synthesize the protein gene products [17]. Both mRNA and protein are subject to degradation and dilution [18]. (See Fig. 3A.) At the single cell level, each of these processes are stochastic. We will model these processes with the stochastic kinetic scheme [17]:

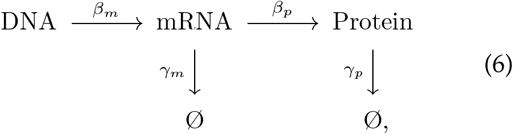

where *β*_*m*_ is the transcription rate (s^*-*1^), *β*_*p*_ is the translation rate (s^*-*1^), *γ*_*m*_ is the message degradation rate (s^*-*1^), and *γ*_*p*_ is the protein effective degradation rate (s^*-*1^). The message lifetime is 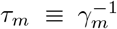. For most protein in the context of rapid growth, dilution is the dominant mechanism of protein depletion and therefore *γ*_*p*_ is approximately the growth rate [15, 19, 20]: *γ*_*p*_ = *T* ^*-*1^ ln 2, where T is the doubling time. We will discuss a more general scenario below.

**FIG. 3.**
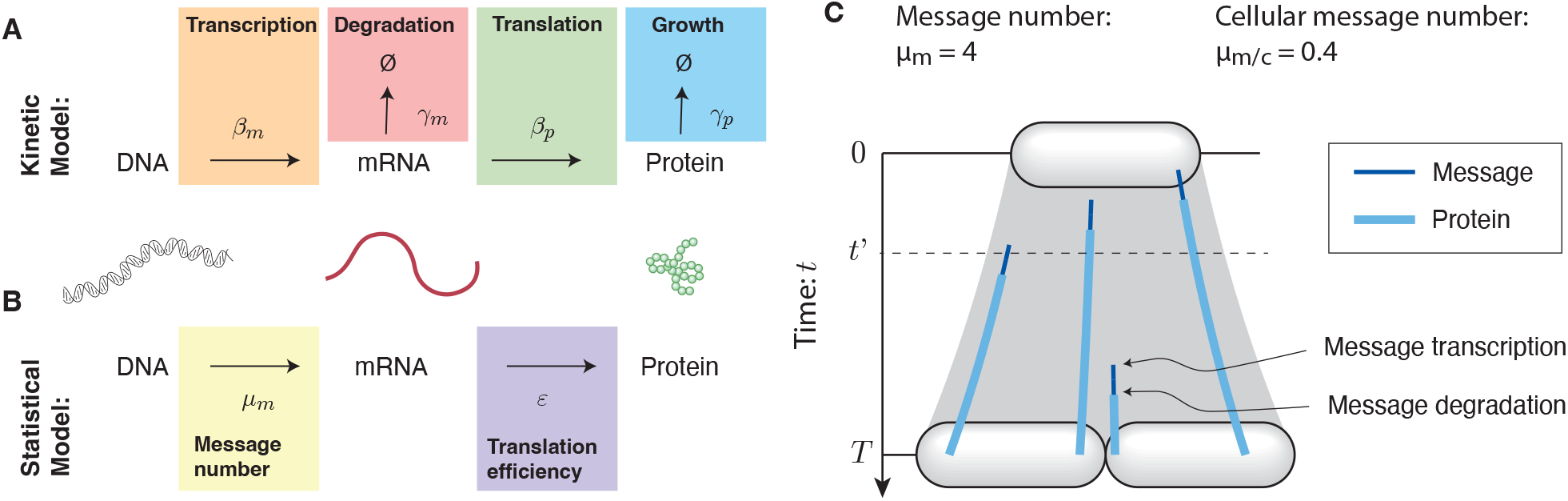
**Panel A: Kinetic model for the central dogma.** The telegraph model is a stochastic kinetic model for protein synthesis, described by four gene-specific rate constants: the transcription rate (*β*_*m*_), the message degradation rate (*γ*_*m*_), the translation rate (*β*_*p*_), and the dilution rate (*γ*_*p*_). **Panel B: Statistical model for the central dogma**. The predicted distribution in protein abundance is described by a gamma distribution, which is parameterized by two unitless constants: the shape parameter *μ*_*m*_, the mean number of messages transcribed per cell cycle, and the scale parameter *ε*, the mean number of proteins translated per message. **Panel C: Message number**. The *message number* (*μ*_*m*_) is defined as the mean total number of messages (dark blue) transcribed per cell cycle. Here, four total messages are transcribed and translated to protein (light blue); however, due to message degradation, at time *t*^*′*^, only one message is present in the cell. Cellular message number (*μ*_*m/c*_) is defined as the mean number of messages per cell at time *t*.

**FIG. 4.**
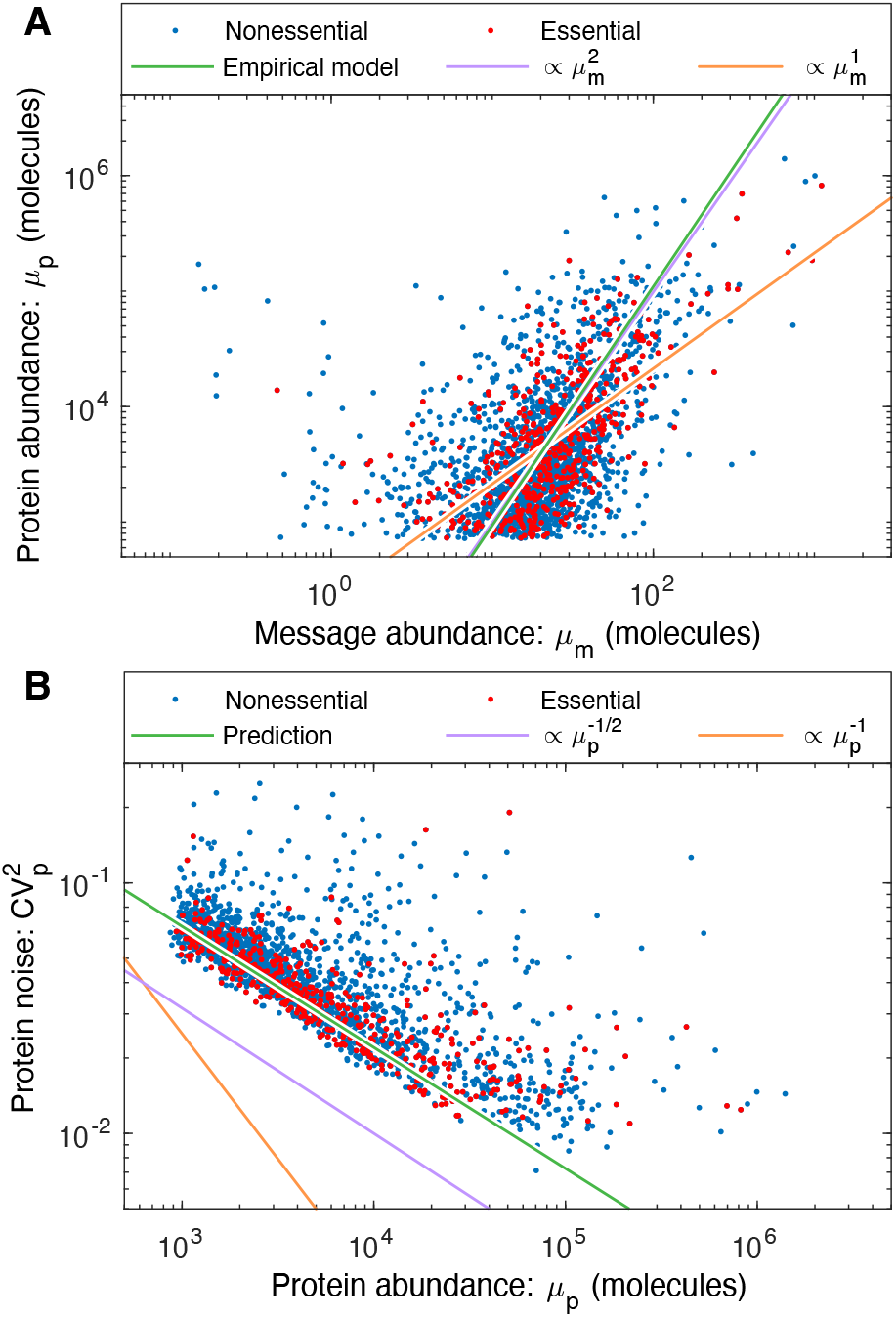
**Panel A: An empirical model for protein number *μ*_*p*_ in yeast.** The canonical noise model assumes constant translation efficiency, which would imply that protein number is proportional to the message number (orange); however, the empirical fit (green) shows that protein number scales close to the square of message number (purple): 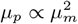. The protein abundance has a cutoff near 10^1^ due to the autofluorescence cutoff [12]. **Panel B: The statistical noise model predicts the observed noise**. The statistical noise model (Eq. 12) and empirical model for protein number (Eq. 15) make a parameter-free prediction of the noise (green). This prediction both closely matches the observed scaling (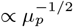, purple) relative to the canonical scaling (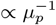, orange) and quantitatively estimates magnitude (vertical offset). This prediction does not include the contribution of noise floor, relevant for describing high-expression proteins.

### Statistical model for protein abundance

To study the stochastic dynamics of gene expression, we used a stochastic Gillespie simulation [21, 22]. (See Supplemental Material Sec. A 1.) In particular, we were interested in the explicit relation between the kinetic parameters (*β*_*m*_, *γ*_*m*_, *β*_*p*_, *γ*_*p*_) and experimental observables.

Consistent with previous reports [3, 9], we find that the distribution of protein number per cell (at cell birth) was described by a gamma distribution: *N*_*p*_ ∼ Γ(*θ*_Γ_, *k*_Γ_), where *N*_*p*_ is the protein number at cell birth and Γ is the gamma distribution which is parameterized by a scale parameter *θ* _Γ_ and a shape parameter *k*_Γ_. (See Supplementary Material Sec. A 1.) The relation between the four kinetic parameters and these two statistical parameters has already been reported, and have clear biological interpretations [9]: The scale parameter:

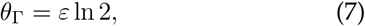

is proportional to the translation efficiency:

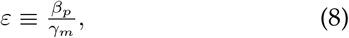

where *β*_*p*_ is the translation rate and *γ*_*m*_ is the message degradation rate. *ε* is understood as the mean number of proteins translated from each message transcribed. The shape parameter *k*_Γ_ can also be expressed in terms of the kinetic parameters [9]:

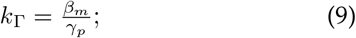

however, we will find it more convenient to express the scale parameter in terms of the cell-cycle message number:

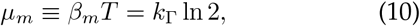

which can be interpreted as the mean number of messages transcribed per cell cycle. Forthwith, we will abbreviate this quantity *message number* in the interest of brevity.

In terms of two gamma parameters, the mean and the squared coefficient of variation are:

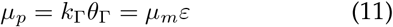

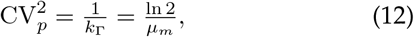

where the noise depends on the message number (*μ*_*m*_), not the mean protein number (*μ*_*p*_). (Eq. 12 only applies when *ε* ≫ 0 [3, 9].) Are these theoretical results consistent with the canonical model (Eq. 3)? We can rewrite the noise in terms of the protein abundance and translation efficiency:

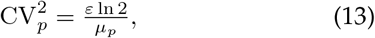

which implies that the canonical model only applies when the translation efficiency (*ε*) is independent of expression (*μ*_*p*_).

### Measuring the message number

The prediction for the noise (Eq. 12) depends on the message number (*μ*_*m*_). However, mRNA abundance is typically characterized by a closely related, but distinct quantity: Quantitative RNA-Seq and methods that visualize fluorescently-labeled mRNA molecules typically measure the number of messages per cell [6]. We will call the mean of this number the *cellular message number μ*_*m/c*_. In the kinetic model, these different message abundances are related:

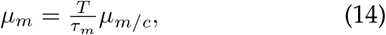

by the message recycling ratio, *T*/*τ*_*m*_, which can be interpreted as the average number of times messages are recycled during the cell cycle. To estimate the message number, we will scale the observed cellular message number *μ*_*m/c*_ by the message recycling ratio, using the mean message lifetime. Fig. 3C illustrates the difference between the message number and the cellular message number. The mean lifetimes, message recycling ratios, as well as the total message number for three model organisms are shown in Tab. I.

**TABLE I.** Central dogma parameters for three model organisms. Columns three through seven hold representative values for measured central-dogma parameters for the model organisms described in the paper. The sources of the numbers and estimates are described in the Supplemental Material Sec. A 2.

| Model organism | Growth condition | Doubling time: $T$ | Message lifetime: $\tau_m$ | Message recycling ratio: $T/\tau_m$ | Total number of | | | Average | |
| --- | --- | --- | --- | --- | --- | --- | --- | --- | --- |
| | | | | | messages /cell: $N_{m/c}^{tot}$ | messages /cell-cycle: $N_m^{tot}$ | proteins: $N_p^{tot}$ | translation efficiency: $\epsilon$ | translation rate: $\beta_p$ ($h^{-1}$) |
| *Escherichia coli* (*E. coli*) | LB | 30 min | 2.5 min | 12 | $7.8 \times 10^3$ | $9.4 \times 10^4$ | $3 \times 10^6$ | 32 | 770 |
| | M9 | 90 min | 2.5 min | 36 | $2.4 \times 10^3$ | $8.6 \times 10^4$ | $3 \times 10^6$ | 35 | 833 |
| *Saccharomyces cerevisiae* (Yeast-haploid) | YEPD | 90 min | 22 min | 4 | $2.9 \times 10^4$ | $1.2 \times 10^5$ | $5 \times 10^7$ | 420 | 1100 |
| *Homo sapiens* (Human) | Tissue | 24 h | 14 h | 1.7 | $3.6 \times 10^5$ | $6.2 \times 10^5$ | $2 \times 10^9$ | $3.2 \times 10^3$ | 230 |

### Construction of an empirical model for protein number

To model the noise as a function of protein abundance (*μ*_*p*_), we will determine the empirical relation between mean protein levels and message abundance by fitting to Eq. 11. Note that the objective here is only to estimate *μ*_*m*_ from *μ*_*p*_, not to model the process mechanistically (*e*.*g*. [23].) The message numbers are estimated from RNA-Seq measurements, scaled as described above (Eq. 14). The protein abundance numbers come from fluorescence and mass-spectrometry based assays [12, 24], with overall normalization chosen to match reported total cellular protein content. (See Supplemental Material Sec. A3 d.) The resulting fit generates our empirical translation model for yeast:

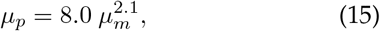

where both means are in units of molecules. (An error analysis for both model parameters is described in Supplementary Material Sec. A4 b.) The data and model are shown in Fig. 5A.

**FIG. 5.**
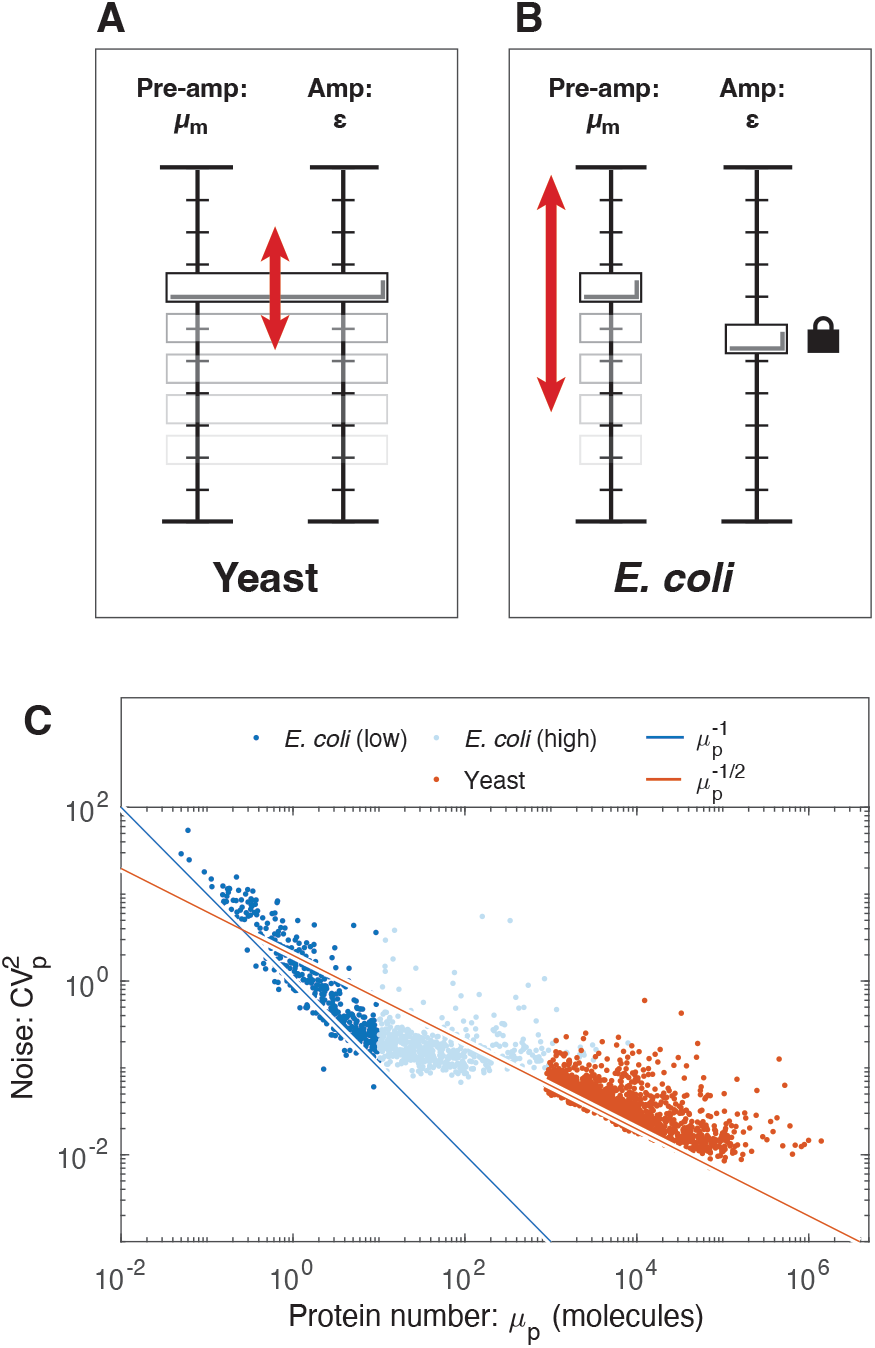
Understanding the distinct central dogma strategies using the amplifier analogy. **Panel A: Yeast.** High expression (*μ*_*p*_) is typically achieved by coordinated small increases in both transcription (*μ*_*m*_) and translation (*ε*), relative to low-expression genes. **Panel B: *E. coli***. High expression (*μ*_*p*_) is typically achieved by a large increase in transcription (*μ*_*m*_) only, relative to low-expression genes. Translation (*ε*) is uncorrelated. **Panel C: Distinct noise scaling with gene expression**. Due to the coordinated changes in both transcription and translation in yeast, noise scaling is weaker than in *E. coli*, where only transcription changes. The noise of high-expression *E. coli* genes is determined by the noise floor.

### Prediction of the noise scaling with abundance

Now that we have fit an empirical model that relates *μ*_*p*_ and *μ*_*m*_, we return to the problem of predicting the yeast noise. We apply the relation (Eq. 15) to Eq. 12 to make a parameter-free prediction of the noise as a function of protein abundance:

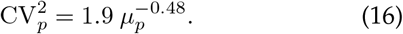

An error analysis for both model parameters is described in Supplementary Material Sec. A4 e. Our noise model (Eq. 16) makes both a qualitative and quantitative prediction: (i) From a qualitative perspective, the model suggests that the *μ*_p_ exponent should be roughly 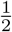 for yeast, rather than the canonically assumed scaling exponent of 1. (ii) From a quantitative perspective, the model also predicts the coefficient of proportionality if the empirical relation between protein and message abundances is known (Eq. 15).

### Observed noise in yeast matches the predictions of the empirical model

Newman *et al*. have characterized protein noise by flow cytometry of strains expressing fluorescent fusions expressed from their endogenous promoters [12]. The comparison of this data to the prediction of the statistical expression model (Eq. 16) are shown in Fig. 5. From a qualitative perspective, the predicted scaling exponent of *-*0.48 comes very close to capturing the scaling of the noise, as determined by the direct fitting of the empirical noise model (Eq. 5 and Fig. 2). From a quantitative perspective, the predicted coefficient of Eq. 16 also fits the observed noise.

From both the statistical analysis (Eq. 5) and visual inspection (Fig. 5C), it is clear that the noise in yeast does not obey the canonical model (Eq. 3). However, the noise in *E. coli* does obey the canonical model for low copy messages [15]. (See Fig. 5C.) Why does the noise scale differently in the two organisms? The key difference is that the empirical relation between the protein and message numbers are different. In *E. coli*, 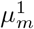 [25]. Our analysis therefore predicts the canonical model (Eq. 3) should hold for *E. coli*, but not for yeast, as illustrated schematically in Fig. 5. (Additional discussion can be found in the Supplementary Material Sec. A5 f.).

### Implications of growth robustness for translation

Before continuing with the noise analysis, we to focus on the significance of the empirical relationship between the protein and message numbers (Eq. 15). How can the cell counteract noise-induced reductions in robustness? Eq. 11 implies that gene expression can be thought of as a two-stage amplifier [17]: The first stage corresponds to transcription with a gain of message number *μ*_*m*_, and the second stage corresponds to translation with a gain in translation efficiency *ε*. (See Fig. 5AB.) The noise is completely determined by the first stage of amplification, provided that *ε* ≫ 0 [3, 9]. Genes with low transcription levels are the noisiest. For these genes, the cell can achieve the same mean gene expression (*μ*_*p*_) with lower noise by increasing the gain of the first stage (increasing message number) and decreasing the gain of the second stage (the translation efficiency) by the same factor. This is most clearly understood by reducing *ε* at fixed *μ*_*p*_ in Eq. 13. Highly transcribed genes have low noise and can therefore tolerate higher translation efficiency in the interest of economy (decreasing the total number of messages) [5]. Growth robustness therefore predicts that the translation efficiency should grow with transcription level.

### Translation efficiency increases with expression level in yeast

The translation efficiency (Eq. 8) can be determined from the empirical translation model (Eq. 15):

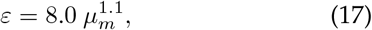

as a function of message number. (An error analysis for both model parameters is described in Supplementary Material Sec. A4 d.) In yeast, the translation efficiency clearly has a strong dependence on message number *μ*_m_, and grows with the expression level, exactly as predicted by robustness arguments. We note the contrast to the translation efficiency in *E. coli*, which is roughly constant [25]. (See Supplementary Material Sec. A5 f.) We will speculate about the rationale for these differences in the discussion below.

### Implications of growth robustness for transcription

In addition to the prediction of translation efficiency depending on transcription, a second qualitative prediction of growth robustness is that essential gene expression should have a noise ceiling, or maximum noise level (Eq. 2), where noise above this level would be too great for robust growth. The fit between the statistical model and the observed noise has an important implication beyond confirming the predictions of the telegraph and statistical models for noise: The identification of the message number, *μ*_*m*_, as the key determinant of noise allows us to use this quantity as a proxy for noise in quantitative transcriptome analysis.

To identify a putative transcriptional floor, we now broaden our consideration beyond yeast to characterize the central dogma in two other model organisms: the bacterium *Escherichia coli* and *Homo sapiens* (human). We will also analyze three different transcriptional statistics for each gene: transcription rate (*β*_*m*_), cellular message number (*μ*_*m/c*_), and message number (*μ*_*m*_). Analysis of these organisms explores orders-of-magnitude differences in characteristics of the central dogma, including total message number, protein number, doubling time, message lifetime, and number of essential genes. (See Tab. I.) In particular, as a consequence of these differences, the three statistics describing transcription: transcription rate, cellular message number and message number are all distinct. Genes with matching message numbers in two different organisms will not have matching transcription rates or cellular message numbers. We hypothesize that cells must express essential genes above some threshold message number for robust growth; however, we expect to see that nonessential genes can be expressed at much lower levels since growth is not strictly dependent on their expression. The signature of a noise-robustness mechanism would be the absence of essential genes for low message numbers.

### No organism-independent threshold is observed for transcription rate or cellular message number

Histograms of the per-gene transcription rate and cellular message number are shown in Fig. 6 for *E. coli*, yeast, and human. Consistent with existing reports, essential genes have higher expression than non-essential genes on average; however, there does not appear to be any consistent threshold in *E. coli* (even between growth conditions), yeast, or human transcription, either as characterized by the transcription rate (*β*_*m*_) or the cellular message number (*μ*_*m/c*_). For instance, the per gene rate of transcription is much lower in human cells than *E. coli* under rapid growth conditions, with yeast falling in between.

**FIG. 6.**
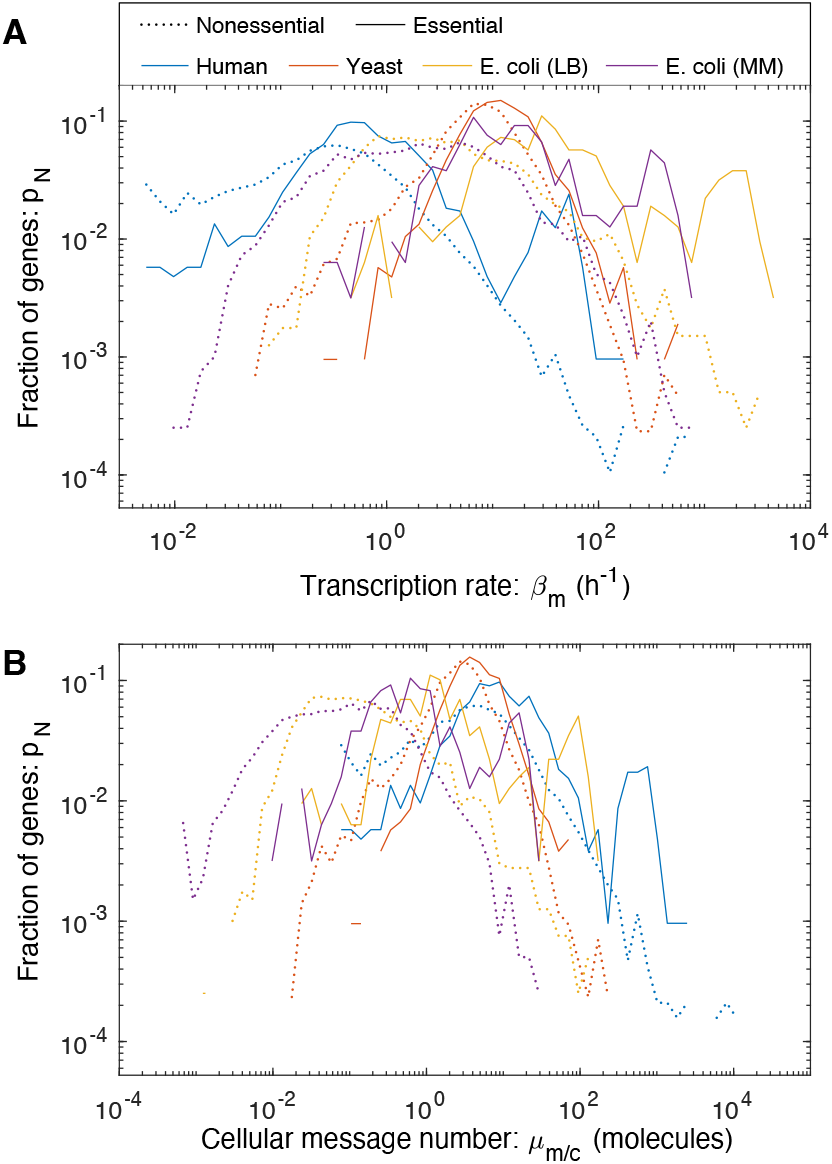
Transcription in three model organisms. We characterized different gene transcriptional statistics in three model organisms. In *E. coli*, two growth conditions were analyzed. **Panel A: The distribution of gene transcription rate**. The transcription rate varies by two orders-of-magnitude between organisms. **Panel B: The distribution of gene cellular message number**. There is also a two-order-of-magnitude variation between cellular message numbers.

### An organism-independent threshold is observed for message number for essential genes

In contrast to the other two transcriptional statistics, there is a consistent lower limit, or floor, on message number (*μ*_*m*_) of some-where between 1 and 10 messages per cell cycle for essential genes. (See Fig. 7.) Non-essential genes can be expressed at a much lower level. This floor is consistent not only between *E. coli*, growing under two different conditions, but also between the three highly-divergent organisms: *E. coli*, yeast and human. We will conservatively define the minimum message number as

**FIG. 7.**
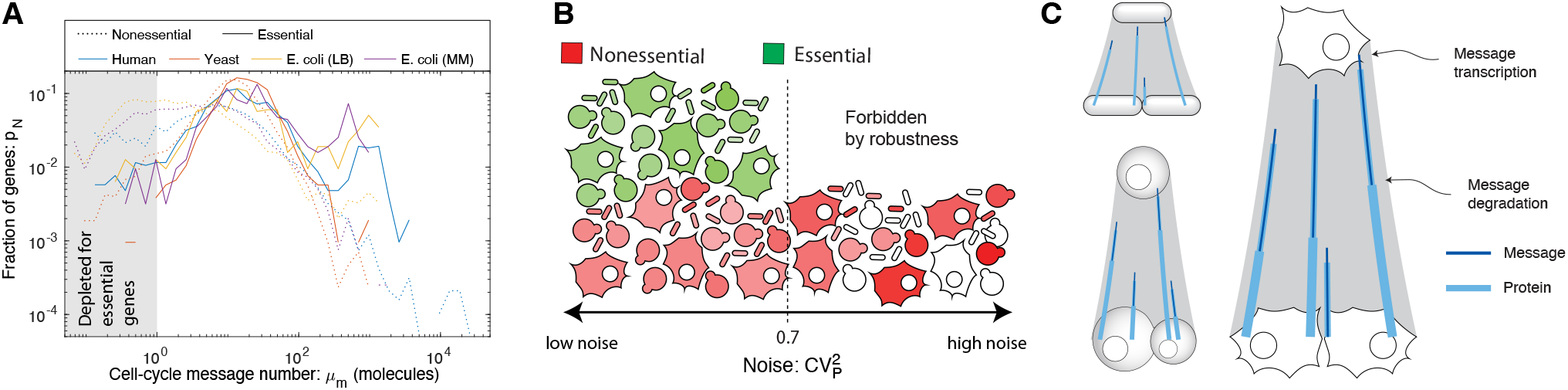
**Panel A: Transcription in three model organisms. The distribution of gene message number.** All organisms have roughly similar distributions of message number for essential genes, which are not observed for message numbers below a couple per cell cycle. However, non-essential genes can be expressed at much lower levels. **Panel B: Nonessential genes tolerate higher noise levels than essential genes**. The floor of message number is consistent with a noise ceiling of 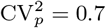 for essential genes (green). Nonessential genes (red) are observed with lower transcription levels. **Panel C: Conserved transcriptional program for essential genes**. The message number per gene (number of messages transcribed per cell cycle) is roughly identical in *E. coli*, yeast, and human. We show this schematically.

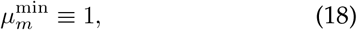

and summarize this observation as the *one-message-per-cell-cycle rule* for essential gene expression.

In addition to the common floor for essential genes, there is a common gene expression distribution shape shared between organisms dependent on the message numbers, especially for low-expression essential genes. This is observed in spite of the significantly larger number of essential genes in human relative to *E. coli*. (See Fig. 7.) Interestingly, there is also a similarity between the non-essential gene distributions for *E. coli* and human, but not for yeast, which appears to have a much lower fraction of genes expressed at the lowest message numbers.

### What genes fall below-threshold?

We have hypothesized that essential genes should be expressed above a threshold value for robustness. It is therefore interesting to consider the function of genes that fall below this proposed threshold. Do functions of these genes give us any insight into essential processes that do not require robust gene expression?

Since our own preferred model system is *E. coli*, we focus here. Our essential gene classification was based on the construction of the Keio knockout library [26]. By this classification, 10 essential genes were below threshold. (See Supplementary Material Tab. IV.) Our first step was to determine what fraction of these genes were also classified as essential using transposon-based mutagenesis [27, 28]. Of the 10 initial candidates, only one gene, *ymfK*, was consistently classified as an essential gene in all three studies, and we estimate that its message number is just below the threshold (μ_m_ = 0.4). *ymfK* is located in the lambdoid prophage element e14 and is annotated as a CI-like repressor which regulates lysis-lysogeny decision [29]. In, \ phase, the CI repressor represses lytic genes to maintain the lysogenic state. A conserved function for *ymfK* is consistent with it being classified as essential, since its regulation would prevent cell lysis. However, since *ymfK* is a prophage gene, not a host gene, it is not clear that its expression should optimize host fitness, potentially at the expense of phage fitness. In summary, closer inspection of below-threshold essential genes supports the threshold hypothesis.

### Maximum noise for essential genes

The motivation for hypothesizing a minimum threshold for message number was noise-robustness, or the existence of a hypothesized noise ceiling above which essential gene expression is too noisy to allow robust cellular proliferation.

With the *one-message-per-cell-cycle rule*, 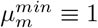, we can estimate the essential gene noise ceiling using Eq. 12:

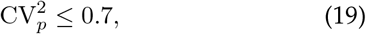

for essential genes. Since noise depends only on the message number, we expect to observe the same limit in all organisms if the message number floor is conserved.

### Estimating the floor on central-dogma parameters

If message number floor is conserved, a limit can be estimated for the floor value on other transcriptional parameters. Using Eq. 14, we can estimate the floor on the cellular message number (as measured in RNA-Seq measurements):

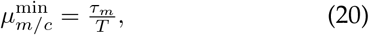

for essential genes. Similarly, we can use Eq. 9 to estimate the minimum transcription rate:

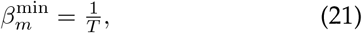

for essential genes. Again, this result has an intuitive interpretation as the one-message-per-cell-cycle rule. Finally, we can estimate a floor on essential protein abundance, assuming a constant translation efficiency using Eq. 11:

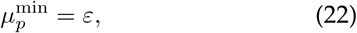

for essential genes, where “ is the translation efficiency (which we will assume is well approximated by the mean in the context of the estimate). All four floor estimates for each model organism are shown in Tab. II.

**TABLE II.** Estimates of threshold levels for the central dogma in three model organisms. Estimates for the lower thresholds of transcription statistics as inferred from our analysis based on the *one-message-per-cell-cycle rule*.

| Model organism | Maximum essential<br>gene noise:<br>$\max CV_p^2$ | Estimated minimum essential gene | | | |
| --- | --- | --- | --- | --- | --- |
| | | messages<br>/cell-cycle:<br>$\mu_m^{\min}$ | messages /cell:<br>$\mu_{m/c}^{\min}$ | transcription rate:<br>$\beta_m^{\min} (h^{-1})$ | proteins:<br>$\mu_p^{\min}$ |
| *E. coli* (LB) | 0.7 | 1 | 0.08 | 2 | 30 |
| (M9) | 0.7 | 1 | 0.03 | 0.7 | 30 |
| Yeast | 0.7 | 1 | 0.2 | 0.7 | 400 |
| Human | 0.7 | 1 | 0.6 | 0.04 | 3000 |

## DISCUSSION

### Noise by the numbers

Although there has already been significant discussion of the scaling of biological noise with protein abundance [3, 9, 10, 12, 15], our study is arguably the first to test the predictions of the telegraph and statistical noise models against absolute measurements of protein and message abundances. This approach is particularly important for the message number (μ_m_), which determines the magnitude of the noise in protein expression, and facilitates direct comparisons of noise between organisms as well as identifying the common distributions of message number for genes, that are conserved from bacteria to human.

### Noise scaling in *E. coli* versus yeast

A key piece of evidence for the significance of the message number was the observation of the non-canonical scaling of the yeast noise with protein abundance (Fig. 5); however, the canonical model (Eq. 3) does accurately describe the noise in *E. coli* (see Fig. 11). Why does the noise scale differently? In *E. coli*, the translation efficiency is only weakly correlated with the gene expression [25], and therefore the canonical model is a reasonable approximation (Supplementary Material Sec. A5 d). However, we also argued that translation efficiency should grow with expression level. Why is this not observed in *E. coli*? Due to the high noise floor in *E. coli*, nearly all essential genes are expressed at a sufficiently high expression level such that the noise is dominated by the noise floor [15]. As a consequence, increasing the message number, while decreasing translation efficiency, does not decrease the noise even as it increases the metabolic load as a result of increased transcription. (A closely related point has recently been made in *Bacillus subtilis* [30], where Deloupy *et al*. report that the noise cannot be tuned by adjusting the message number due to the noise floor.) Our expectation is therefore that other bacterial cells will look similar to *E. coli*: They will have a higher noise floor and a similar scaling of noise with protein abundance.

In contrast, due to the lower noise floor, we expect eukaryotic cells to optimize the central dogma processes like yeast and as a result will have a similar non-canonical scaling of noise with protein abundance. Although this non-canonical scaling is clear from the abundance data (Fig. 5B), there is an important qualification to emphasize: the mechanism that gives rise to the non-canonical scaling is due to the correlation between translation efficiency and transcription. Regulatory changes that effect only transcription (*i*.*e*. increase μ_m_) and not translation (“) should obey the canonical noise model (Eq. 3). This scenario may help explain why Bar-Even *et al*. claim to observe canonical noise scaling in yeast [10], studying a subset of genes under a range of conditions resulting in differential expression levels. The failure of the canonical noise model (Eq. 3) at the proteome level in yeast (Eq. 16) is a consequence of genome-wide optimization of the relative transcription and translation rates.

### Essential versus non-essential genes

What genes are defined as *essential* is highly context specific [31]. It is therefore important to consider whether the comparison between these two classes of genes is informative in the context of our analysis. We believe the example of *lac* operon in *E. coli* is particularly informative in this respect. The genes *lacZYA* are conditionally essential: they are required when lactose is the carbon source; however, these genes are repressed when glucose is the carbon source. Our expectation is that these conditionally essential genes will obey the one-message-per-cell-cycle rule when these genes are required; however, they need not obey this rule when the genes are repressed. By analyzing essential genes, we are limiting the analysis to transcriptionally-active genes, whereas the non-essential category contains both transcriptionally-active and silenced genes.

### Protein degradation and transcriptional bursting

Two important mechanisms can act to significantly increase the noise above the levels we predict: protein degradation and transcriptional bursting. Although the dominant mechanism of protein depletion is dilution in *E. coli*, protein degradation plays an important role in many organisms, especially in eukaryotic cells [32, 33]. If protein degradation depletes proteins faster than dilution, the shape parameter decreases below our estimate (Eq. 9), increasing the noise. Likewise, the existence of transcriptional bursting, in which the chromatin switches between transcriptionally active and quiescent periods, can also act to increase the noise [1, 7, 34]. Since the presence of both these mechanisms increases the noise beyond what is predicted by the message number, they do not affect our estimate of the minimum threshold for μ_m_.

### The biological implications of noise

What are the biological implications of gene expression noise? Many important proposals have been made, including bethedging strategies, the necessity of feedback in gene regulatory networks, *etc* [1]. Our analysis suggests that noise influences the optimal function of the central dogma process generically. Hausser *et al*. have already discussed some aspects of this problem and use this approach to place coarse limits on transcription versus translation rates [5]. The transcriptional floor for essential genes that we have proposed places much stronger limits on the function of the central dogma.

Although we describe our observations as a floor, a more nuanced description of the phenomenon is a common distribution of gene message numbers, peaked at roughly 15 messages per cell cycle and cutting off close to one message per cell cycle. Does this correspond to a hard limit? We expect that this does not since there are a small fraction of genes, classified as essential, just below this limit; however, it does appear that virtually all essential genes have optimal expression levels above this threshold. The common distribution of message number clearly suggests that noise considerations shape the function of the central dogma for virtually all genes. Exploring this hypothesis will require quantitative models that explicitly realize the high cost of noise-induced low essential-protein abundance. We will present such an analysis elsewhere.

### Adapting the central dogma to increased cell size and complexity

Although core components of the central dogma machinery are highly-conserved, there has been significant complexification of both the transcriptional and translational processes in eukaryotic cells [35]. Given this increased regulatory complexity, it is unclear how the central dogma processes should be adapted in larger and more complex cells. An important clue to this adaptation comes from *E. coli* proliferating with different growth rates. Although there are very significant differences between the cellular message number as well as the overall transcription rate under the two growth conditions, there is very little difference in message number. In short, roughly the same number of messages are made during the cell cycle, but they are made more slowly under slow growth conditions.

How does this picture generalize in eukaryotic cells? Although both the total number of messages and the number of essential and non-essential genes are larger in both yeast and human cells, the distribution of the message number per gene is essentially the same as *E. coli* (Fig. 7). The conservation of the message number between organisms is consistent with all of these organisms being optimized with respect to the same trade-off between economy and robustness to noise.

## Data availability

We include a source data file which includes the estimated message numbers as well as essential/nonessential classifications for each organism.

## Acknowledgments

The authors would like to thank B. Traxler, A. Nourmohammad, J. Mougous, K. Cutler, M. Cosentino-Lagomarsino, S. van Teeffelen, and S. Murray. This work was supported by NIH grant R01-GM128191.

## Author contributions

T.W.L., H.K.J.C., D.H. and P.A.W. conceived the research. T.W.L. and P.A.W. performed the analysis. H.K.J.C. and D.H. performed experiments and analysis. T.W.L., H.K.J.C., D.H. and P.A.W. wrote the paper.

