## Supplemental Material for "The one-message-per-cell-cycle rule: A conserved minimum transcription level for essential genes"

**CONTENTS**

Introduction 1

Results 2

Discussion 8

References 10

A. Supplemental analysis 12
1. Gillespie Simulation of the telegraph model 12
2. Selection of central dogma parameter estimates 12
a. *E. coli* data 13
b. Yeast data 13
c. Human data 14
3. Quantitative estimates of central dogma parameters 14
a. Estimating the *cellular message number*: $\mu_{m/c}$ 14
b. Estimating the *transcription rate*: $\beta_m$ 14
c. Estimating the *message number*: $\mu_m$ 14
d. Estimating the *protein number*: $\mu_p$ 14
4. Empirical models for yeast gene expression 15
a. The meaning of the error estimates 15
b. Empirical model for protein number 15
c. Empirical model for message number 15
d. Empirical model for translation efficiency 15
e. Empirical model for the coefficient of variation 16
5. Supplemental analysis of gene expression noise 16
a. Inclusion of noise floor in the yeast analysis 16
b. Hypothesis test I 16
c. Hypothesis test II 17
d. Maximum likelihood estimates of the parameters 17
e. Statistical details MLE estimate of the variance 17
f. Detailed discussion of noise in *E. coli* 17

### Appendix A: Supplemental analysis

#### 1. Gillespie Simulation of the telegraph model

Protein distributions of the telegraph model for *E. coli* were simulated with a Gillespie algorithm. Assuming the lifetime of the cell cycle ($T_{cc} = 30$ min) [36], mRNA lifetime ($\tau_m = 2.5$ min) [37], and translation rate ($\beta_p \approx 500$ hr$^{-1}$), the protein distributions for several mean expression levels were numerically generated for exponential growth with 100,000 stochastic cell divisions, with protein partitioned at division following the binomial distribution.

The gamma distributions for each mean message number with scale and shape parameters determined by the corresponding translation efficiency and message number ($\theta = \varepsilon \ln 2$, $k = \frac{\mu_m}{\ln 2}$) as used for the Gillespie simulation were also plotted with the protein distributions.

$$p(n|\theta, k) = \frac{1}{\Gamma(k)\theta^k} n^{k-1} e^{-\frac{n}{\theta}} \quad \text{(A1)}$$

#### 2. Selection of central dogma parameter estimates

The estimates for central dogma model parameters come from two types of data: (i) quantitative measurement of cellular-scale parameters for each organism (total number of messages in the cell, cell cycle duration, *etc*) and (ii) genome-wide studies quantitative of mRNA and protein abundance.

For the cellular-scale central dogma parameters, we relied heavily on an online compilation of biological numbers: BioNumbers [45]. This resource provides a collection of curated quantitative estimates for biological numbers, as well as their original source. In the interest of conciseness, we have cited only the original source in the Tab. I, although we are extremely grateful and supportive of the creators of the BioNumbers website for helping us very efficiently identify consensus estimates for the parameters of the central dogma parameters.

For the selection of genome-wide studies on abundance, we used many of the same resources cited in BioNumbers as well as studies selected by a previous study of a quantitative analysis of the central dogma: Hausser *et al.* [5].

| Model organism | Growth condition | Doubling time: $T$ | Message lifetime: $\tau_m = \gamma_m^{-1}$ | Message recycling ratio: $m = T/\tau_m$ | Total number of messages /cell: $N_{m/c}^{\text{tot}}$ | Total number of messages /cell-cycle: $N_m^{\text{tot}}$ | Total number of proteins: $N_p^{\text{tot}}$ | Average translation efficiency: $\varepsilon$ | Average translation rate: $\beta_p$ (h$^{-1}$) |
|---|---|---|---|---|---|---|---|---|---|
| *Escherichia coli (E. coli)* | LB | 30 min [36] | 2.5 min [37] | 12 | $7.8 \times 10^3$ [38] | $9.4 \times 10^4$ | $3 \times 10^6$ [39] | 22 | 530 |
| | M9 | 90 min [36] | 2.5 min [37] | 36 | $2.4 \times 10^3$ [38] | $8.6 \times 10^4$ | $3 \times 10^6$ [39] | 24 | 580 |
| *Sacchromyces cerevisiae (Yeast–haploid)* | YEPD | 90 min [40] | 22 min [41] | 4 | $2.9 \times 10^4$ [42] | $1 \times 10^5$ | $5 \times 10^7$ [43] | $4 \times 10^2$ | 410 |
| *Homo sapiens (Human)* | Tissue | 24 h [35] | 14 h [44] | 1.7 | $3.6 \times 10^5$ [40] | $5 \times 10^5$ | $2 \times 10^9$ [39] | $4 \times 10^3$ | 120 |

TABLE III. **Central dogma parameters for three model organisms with detailed references.** Columns three through seven hold representative values for measured central-dogma parameters for the model organisms described in the paper. Each value is followed by a reference for its source. The last three columns hold estimates for the lower threshold on transcription inferred from our analysis. The sources of the numbers and estimates are described in the Supplemental Material Sec. A 2.

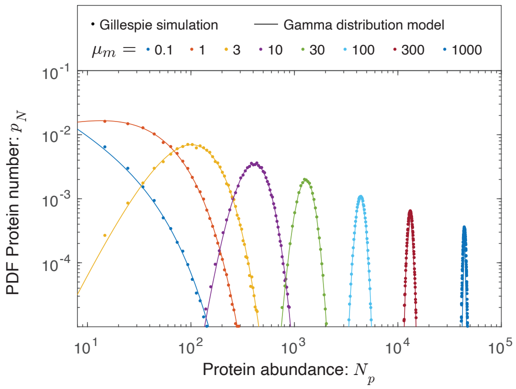


FIG. 8. **The protein abundance is approximately gamma distributed.** Protein abundance was modeled for eight different transcription rates using a Gillespie simulation, including the stochastic partitioning of the proteins between daughter cells at cell division. The range in abundance matches the observed range of expression levels in the cell. We observed that the simulated protein abundances were well fit by gamma distributions.

##### a. *E. coli data*

**Message lifetimes:** The message lifetimes (and median lifetime) were taken from a recent transcriptome-wide study by Chen *et al.* [37]. These investigators measured the lifetime in both rapid (LB) and slow growth (M9).

**Noise:** Taniguchi *et al.* have performed a beautiful simultaneous study of the proteome and transcriptome with single-molecule sensitivity [15]. Although we use the noise analysis data from this study for our supplemental analysis of *E. coli* noise, it is not the source for our *E. coli* transcriptome data due to the extremely slow growth of the cells in this study (150 minute doubling time), which is not consistent with the growth conditions for the other sources of data.

**mRNA abundance:** Instead, we used data from the more recent Bartholomaus *et al.* study [38], which characterizes the transcriptome in both rapid (LB) and slow growth (M9).

**Total cellular message number.** This study was chosen since it was the source of the BioNumbers estimates of cellular message number in *E. coli* (BNID 112795 [45]).

**Doubling time:** The source of the doubling times for rapid (LB) and slow (M9) growth of *E. coli* comes from Bernstein [36].

**Essential gene classification.** The classification of essential genes in yeast comes from the construction of the Keio knockout collection from Baba *et al.* [26].

**Protein number.** The total protein number in *E. coli* came from Milo's recent review of this subject [39].

##### b. *Yeast data*

**Message lifetimes:** The message lifetimes (and median lifetime) were taken from Chia *et al.* [41].

**Noise:** The noise data was taken from the Newman *et al.* study, which used flow cytometry of a library of fluorescent fusions to characterize protein abundance with single-cell resolution [12].

**mRNA abundance:** The transcriptome data comes from the very recent Blevins *et al.* study [46].

**Total cellular message number.** There are a wide-range of estimates for the total cellular message number in yeast: $1.5 \times 10^4$ [47] (BNID 104312 [45]), $1.2 \times 10^4$ [48] (BNID 102988 [45]), $6.0 \times 10^4$ [49] (BNID 103023 [45]), $2.6 \times 10^4$ [42] (BNID 106763 [45]) and $3.0 \times 10^4$ [50]. We used the compromise value of $2.9 \times 10^4$.

**Doubling time:** The doubling time was taken from [40].

**Protein number.** The total protein number in yeast comes from Futcher *et al.* [43].

**Essential gene classification.** The classification of essential genes in yeast comes from van Leeuwen *et al.* [51].

**Proteome abundance data:** The proteome abundance data came from two sources: flow cytometry of fluorescent fusions from Newman *et al.* [12] as well as mass-spec data from de Godoy *et al.* [24].

##### c. Human data

**Message lifetimes:** The message lifetimes (and median lifetime) were taken from Yang *et al.* [44] who reported a median half life of 10 h which corresponds to a lifetime of 14 h.

**mRNA abundance:** The transcriptome data comes from the data compiled by the Human Protein Atlas [52], which we averaged over tissue types.

**Total cellular message number.** The total cellular message number in human comes from Velculescu *et al.* [53] (BNID 104330 [45]).

**Doubling time:** The doubling time was taken from [35].

**Protein number.** The total protein number in human came from Milo's recent review of this subject [39].

**Essential gene classification.** The classification of essential genes in human comes from Wang *et al.* [54].

#### 3. Quantitative estimates of central dogma parameters

##### a. Estimating the cellular message number: $\mu_{m/c}$

For each model organism (and condition), we found a consensus estimate from the literature for the total number of mRNA messages per cell $N^{\text{tot}}_{m/c}$. This number and its source are provided in Tab. I. To estimate the number of messages corresponding to gene $i$, we re-scaled the un-normalized abundance level $r_i$:

$$N_{m/c,i} = N^{\text{tot}}_{m/c}\frac{r_i}{\sum_j r_j}, \tag{A2}$$

where the sum over gene index $j$ runs over all genes.

##### b. Estimating the transcription rate: $\beta_m$

To estimate the transcription rate for gene $i$, we start from the estimated cellular message number $N_{m/c,i}$ and use the telegraph model prediction for the cellular message number:

$$N_{m/c,i} = \beta_{m,i}/\gamma_{m,i}, \tag{A3}$$

where $\gamma_{m,i}$ is the message decay rate. Since gene-to-gene variation in message number is dominated by the transcription rate (*e.g* [37]), we estimate the decay rate as the inverse gene-median message life time:

$$\gamma_{m,i} = \tau_m^{-1}, \tag{A4}$$

for which a consensus value was found from the literature. This number and its source are provided in Tab. I. We then estimate the gene-specific transcription rate:

$$\beta_{m,i} = N_{m/c,i}/\tau_m. \tag{A5}$$

##### c. Estimating the message number: $\mu_m$

To estimate the message number of gene $i$, we use the predicted value from the telegraph model:

$$N_{m,i} = T\beta_{m,i} = \tfrac{T}{\tau_m}N_{m/c,i}, \tag{A6}$$

where $T$ is the doubling time and $N_{m/c,i}$ is the cellular message number (Eq. A2).

##### d. Estimating the protein number: $\mu_p$

The protein abundance data for yeast grown in YEPD media and measured with flow cytometry fluorescence [12] were given in arbitrary units (AU). In order to convert from AU to protein number, the fluorescence values were rescaled by comparing with mass-spectrometry protein abundance data for yeast grown in YNB media [24]. Since the protein abundance from mass-spectrometry was given in terms of Intensity, the Intensity values were first rescaled by the total number of proteins in yeast, $5 \times 10^7$. The mass-spectrometry protein data was thresholded at 10 proteins, based on the assumption that the noise of the data for 10 and fewer proteins makes the data unreliable. Next, the log of the fluorescence protein abundance in AU as a function of the log of thresholded mass-spectrometry protein abundance was fit as a linear function with an assumed slope of 1 to find the offset, 3.9, (Fig. 9) which corresponds to a multiplicative scaling factor (Eqn. A8). We then used that offset value to rescale the fluorescence data from AU to protein number. We also compared to yeast grown in SD media [12] and found a similar offset result.

$$\log \mu_P^{\text{F}} = m \log \mu_P^{\text{MS}} + b \tag{A7}$$

$$\mu_P^{\text{F}} = b(\mu_P^{\text{MS}})^m \tag{A8}$$

#### 4. Empirical models for yeast gene expression

To generate the empirical model for protein number as a function of message number, we used protein abundance data from Newman *et al.* [12], re-scaled to estimate protein number (Sec. A 3 d) and transcriptome

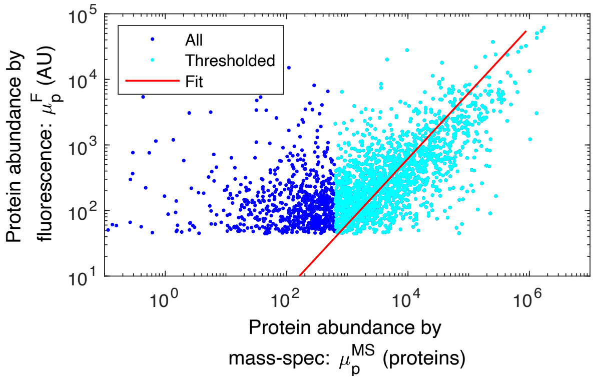


FIG. 9. **Fit to rescale fluorescence intensity to protein number.** Protein abundance from flow cytometry fluorescence [12] as a function of mass-spectrometry scaled abundance [24]. The mass-spectrometry data was thresholded at 10 proteins, and then a linear fit was performed to find the offset of 3.9, which was used to convert protein fluorescence AU to number.

data from Lahtvee *et al.* [55], re-scaled to estimate message number (Sec. A 3 c).

##### a. *The meaning of the error estimates*

Before providing a detailed error analysis, it is important to place our error estimates in perspective. The error that we will be estimating is the statistical error associated with the finite sample size; however, *this is not the dominant source of error*. A far more important consideration are systematic problems with our analysis and the underlying experiments. For instance, since we do not have a detailed model for the error of the experiments analyzed, there are multiple distinct analyses (*i.e.* assumptions about the error model) that could be implemented for the data fitting, each giving slightly different model parameters. These model to model differences still give rise to predictions consistent with our qualitative conclusions; however, they are likely larger than the statistical uncertainty we compute (while assuming a particular model).

##### b. *Empirical model for protein number*

We initially fit the empirical model for protein number,

$$\mu_p = C_0 \mu_m^{\alpha_0}, \tag{A9}$$

to the data using a standard least-squares approach; however, the algorithm led to a very poor fit since it does not account for uncertainty in both independent and dependent variables. We therefore used an alternative approach [56], which assumes comparable error in both variables. The model parameters are:

$$\alpha_0 = 2.1 \pm 0.04, \tag{A10}$$
$$C_0 = 8.0 \pm 1.0, \tag{A11}$$

where the uncertainties are the estimated standard errors.

##### c. *Empirical model for message number*

For the prediction of the coefficient of variation, it is useful to invert Eq. A9 to generate a model for message number as a function of protein number:

$$\mu_m = C_0^{-1/\alpha_0} \mu_p^{1/\alpha 0}, \tag{A12}$$
$$= C_1 \mu_p^{\alpha_1}, \tag{A13}$$

where the last line defines two new parameters: a coefficient $C_1$ and an exponent $\alpha_1$. The resulting parameters and uncertainties are:

$$\alpha_1 \equiv 1/\alpha_0, \tag{A14}$$
$$= 0.48 \pm 0.01, \tag{A15}$$
$$C_1 \equiv C_0^{-1/\alpha_0}, \tag{A16}$$
$$= 0.37 \pm 0.02, \tag{A17}$$

where the uncertainties are the estimated standard errors.

##### d. *Empirical model for translation efficiency*

To generate an empirical model for translation efficiency, we started from the empirical model for protein number (Eq. A9), and then use Eq. 11 to relate protein number, message number, and translation efficiency:

$$\varepsilon = \frac{\mu_p}{\mu_m}, \tag{A18}$$
$$= C_0 \mu_m^{\alpha_0 - 1}, \tag{A19}$$
$$= C_2 \mu_m^{\alpha_2}, \tag{A20}$$

where the last line defines two new parameters: a coefficient $C_2$ and an exponent $\alpha_2$. The resulting parameters and uncertainties are:

$$\alpha_2 = \alpha_0 - 1, \tag{A21}$$
$$= 1.07 \pm 0.04, \tag{A22}$$
$$C_2 = C_0, \tag{A23}$$
$$= 8.0 \pm 1.0, \tag{A24}$$

where the uncertainties are the estimated standard errors.

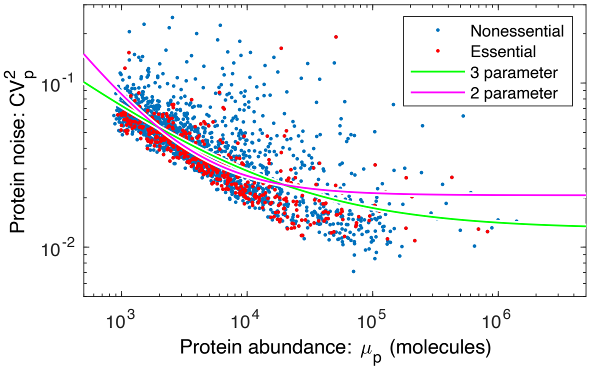


FIG. 10. **Yeast noise fit against canonical noise model, with a noise floor.** Yeast noise data fit with the 2- (null hypothesis with $\mu_p^{-1}$ dependence) and 3- parameter ($\mu_p^a$) models.

*e. Empirical model for the coefficient of variation*

To generate an empirical model for the coefficient of variation, we started from the empirical model for message number (Eq. A13), and then substitute this into the statistical model prediction for $\text{CV}_p^2$ (Eq. 12):

$$\text{CV}_p^2 = \frac{\log 2}{\mu_m}, \tag{A25}$$

$$= C_0^{1/\alpha_0} \log 2 \cdot \mu_p^{-1/\alpha_0}, \tag{A26}$$

$$= C_3 \mu_p^{\alpha_3}, \tag{A27}$$

where the last line defines two new parameters: a coefficient $C_3$ and an exponent $\alpha_3$. The resulting parameters and uncertainties are:

$$\alpha_3 \equiv -1/\alpha_0, \tag{A28}$$

$$= -0.48 \pm 0.01, \tag{A29}$$

$$C_3 \equiv C_0^{1/\alpha_0} \log 2, \tag{A30}$$

$$= 1.9 \pm 0.1, \tag{A31}$$

where the uncertainties are the estimated standard errors.

#### 5. Supplemental analysis of gene expression noise

The quantitative model for gene expression noise includes multiple contributions:

$$\text{CV}_p^2 \approx \frac{1}{\mu_p} + \frac{\log 2}{\mu_m} + c_0, \tag{A32}$$

where the first term can be understood to represent the Poisson noise from translation, the second term the Poisson noise from transcription, and the last term, $c_0$, is called the *noise floor* and is believed to be caused by the cell-to-cell variation in metabolites, ribosomes, and polymerases *etc* [13, 14].

*a. Inclusion of noise floor in the yeast analysis*

In the main text of the paper, we have ignored the role of the noise floor in the analysis of noise in yeast. Unlike *E. coli*, where the noise floor is high ($\text{CV}_p^2 = 0.1$) and is determinative of the noise associated with almost all essential genes [13–15], in yeast the noise floor is much lower ($\text{CV}_p^2 = 0.01$) and therefore affects only genes with the highest expression.

In this section, we will consider models that include the noise floor, since its presence can make the noise scaling more difficult to interpret. To determine if the scaling of the noise is consistent with the canonical assumption that the noise is proportional to $\mu_p^{-1}$ for low expression, we will consider two competing empirical models for the noise (Fig. 10). In the null hypothesis, we will consider a model:

$$\eta_0(\mu_p; b, c) = \frac{b}{\mu_p} + c, \tag{A33}$$

and an alternative hypothesis with an extra exponent parameter $a$:

$$\eta_1(\mu_p; a, b, c) = \frac{b}{\mu_p^a} + c. \tag{A34}$$

We will assume that $\text{CV}_p^2$ is normally distributed about $\eta$ with unknown variance $\sigma_\eta^2$.

In this context, a maximum likelihood analysis is equivalent to least-squares analysis. Let the sum of the squares be defined:

$$S_I(\boldsymbol{\theta}) \equiv \sum_i [\text{CV}_{p,i}^2 - \eta_I(\mu_{p,i}; \boldsymbol{\theta})]^2 \tag{A35}$$

for model $I$ where $\boldsymbol{\theta}$ represents the parameter vector. The maximum likelihood parameters are

$$\hat{\boldsymbol{\theta}} = \arg\max_{\boldsymbol{\theta}} S_I(\boldsymbol{\theta}), \tag{A36}$$

with residual norm:

$$\hat{S}_I = S_I(\hat{\boldsymbol{\theta}}). \tag{A37}$$

To test the null hypothesis, we will use the canonical likelihood ratio test with the test statistic:

$$\Lambda \equiv 2 \log \frac{q_1}{q_0}, \tag{A38}$$

where $q_0$ and $q_1$ are the likelihoods of the null and alternative hypotheses, respectively. Wilks' theorem states that $\Lambda$ has a chi-squared distribution of dimension equal to the difference of the dimension of the alternative and null hypotheses ($3 - 2 = 1$).

*b. Hypothesis test I*

In our first analysis, we will estimate the variance directly. We computed the mean-squared difference for successive $\mathrm{CV}_p^2$ values, sorted by mean protein number $\mu_p$. The variance estimator is

$$\hat{\sigma}_\eta^2 = \tfrac{1}{2}\left\langle (\mathrm{CV}_{p,i}^2 - \mathrm{CV}_{p,i+1}^2)^2 \right\rangle_i = 6.3 \times 10^{-4}, \quad \text{(A39)}$$

where the brackets represent a standard empirical average over gene $i$ for the $\mu_p$-ordered gene $\mathrm{CV}_p^2$ values. The test statistic can now be expressed in terms of the residual norms:

$$\Lambda = (\hat{S}_1 - \hat{S}_2)/\hat{\sigma}_\eta^2, \quad \text{(A40)}$$
$$= 3.3 \times 10^4, \quad \text{(A41)}$$

which corresponds to a p-value far below machine precision. We can therefore reject the null hypothesis.

#### c. *Hypothesis test II*

In a more conservative approach, we can use maximum likelihood estimation to estimate the variance of each model independently as a model parameter. In this case, the test statistic can again be expressed in terms of the residual norms:

$$\Lambda = N \log \tfrac{\hat{S}_1}{\hat{S}_2}, \quad \text{(A42)}$$
$$= 1.6 \times 10^2, \quad \text{(A43)}$$

where $N$ is the number of data points. In this case, the p-value can be computed assuming the Wilks' theorem (*i.e.* the chi-squared test):

$$p = 6 \times 10^{-36}, \quad \text{(A44)}$$

again, strongly rejecting the null hypothesis.

#### d. *Maximum likelihood estimates of the parameters*

In the alternative hypothesis, the maximum likelihood estimate (MLE) of the empirical noise model (Eq. 5) parameters are (Fig. 10):

$$a = 0.57 \pm 0.02, \quad \text{(A45)}$$
$$b = 3.0 \pm 0.5, \quad \text{(A46)}$$
$$c = 0.013 \pm 0.001, \quad \text{(A47)}$$

where the parameter uncertainty has been estimated using the Fisher Information in the usual way using the MLE estimate of the variance.

The noise model parameters were also determined for *E. coli*:

$$a = 1.22 \pm 0.01, \quad \text{(A48)}$$
$$b = 1.27 \pm 0.02, \quad \text{(A49)}$$
$$c = 0.154 \pm 0.002, \quad \text{(A50)}$$

with the corresponding fit shown in Fig. 11. Since $a$ is close to 1, the canonical model with $a = 1$ (Eqn. 3) is a somewhat reasonable approximation for the noise in *E. coli*.

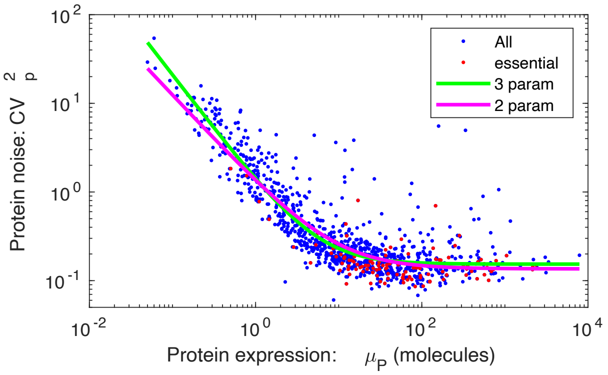


FIG. 11. **Three-parameter fit to *E. coli* noise.** The noise as a function of protein abundance from Taniguichi *et al.* was fit to the 3 parameter noise model (Eqn. 5). From the fit, protein noise scales proportionally with $\mu_p^{-1.22}$, which is a close result to the canonical model with $\mu_p^{-1}$.

#### e. *Statistical details MLE estimate of the variance*

The minus-log-likelihood for the normal model $I$ is:

$$h_I(\hat{\boldsymbol{\theta}}, \sigma^2) = \tfrac{N}{2}\log 2\pi\sigma^2 + \tfrac{1}{2\sigma^2}\hat{S}_I, \quad \text{(A51)}$$

where $\hat{S}_I$ is the least-square residual. We then minimize $h_I$ with respect to the variance $\sigma^2$:

$$\partial_{\sigma^2} h|_{\hat{\sigma}^2} = 0, \quad \text{(A52)}$$

to solve for the MLE $\hat{\sigma}^2$:

$$\hat{\sigma}^2 = \tfrac{1}{N}\hat{S}_I. \quad \text{(A53)}$$

Next we evaluate $h$ at the variance estimator:

$$h_I(\hat{\boldsymbol{\theta}}, \hat{\sigma}^2) = \tfrac{N}{2}\left[\log 2\pi \tfrac{\hat{S}_I}{N} + 1\right]. \quad \text{(A54)}$$

The test statistics can be written in terms of the $h$'s:

$$\Lambda = 2h_0(h_1(\hat{\boldsymbol{\theta}}, \hat{\sigma}^2) - 2h_2(\hat{\boldsymbol{\theta}}, \hat{\sigma}^2), \quad \text{(A55)}$$
$$= N \log \tfrac{\hat{S}_1}{\hat{S}_1}, \quad \text{(A56)}$$

which can be evaluated directly in terms of the residual norms for the null and alternative hypotheses.

#### f. *Detailed discussion of noise in E. coli*

In general, the Telegraph model predicts that the noise will have a coefficient of variation [3, 9]:

$$\mathrm{CV}_p^2 \approx \frac{1}{\mu_p} + \frac{\varepsilon \ln 2}{\mu_p}, \quad \text{(A57)}$$

where the first term is significant whenever the translation efficiency isn't $\varepsilon \gg 1$. In both *E. coli* ($\varepsilon \approx 30$) and yeast ($\varepsilon \approx 420$), this would seem naively to be the case.

| Gene name | Message number: $\mu_m$ | Annotated function from Ecocyc | Essential (E)/ Nonessential (N) Ref. [26], [57], [28] |
|---|---|---|---|
| *alsK* | 0.3 | The alsK gene encodes a D-allose kinase. Its role in the degradation of D-allose is unclear; AlsK is not required for utilization of a D-allose carbon source; this effect may be due to the presence of other ambiguous sugar kinases within *E. coli* K-12. | E, N, N |
| *bcsB* | 0.4 | BcsB is encoded in a predicted operon together with *bcsA*, *bcsZ* and *bcsC*. In other organisms, these genes are involved in cellulose biosynthesis, a characteristic of the rdar (red, dry and rough) morphotype. However, the K-12 laboratory strain of *E. coli* does not show a rdar morphotype and does not produce cellulose. | E, N, N |
| *entD* | 0.4 | AcpS is the founding member of a 4'-phosphopantetheinyl (P-pant) transferase protein family that includes *E. coli* EntD, *E. coli* o195 protein, and *Bacillus subtilis* Sfp; family members share two conserved motifs but relatively low sequence identity overall. | E, N, N |
| *yafF* | 0.4 | No information about this protein was found by a literature search conducted on April 19, 2017. | E,-, N |
| *yagG* | 0.6 | *yagGH* is predicted to be a member of the XylR regulon; its products may mediate transport (YagG) and hydrolysis (YagH) of xylooligosaccharides; putative XylR and CRP binding sites are identified upstream of *yagGH*. | E,-, N |
| *yceQ* | 0.2 | No information about this protein was found by a literature search conducted on July 12, 2017. | E, E, N |
| *ydiL* | 0.2 | No information about this protein was found by a literature search conducted on April 7, 2017. | E, N, N |
| *yhhQ* | 0.4 | YhhQ is an inner membrane protein implicated in the uptake of queuosine (Q) precursors - 7-cyano-7-deazaguanine (preQ0) and 7-aminomethyl-7-deazaguanine (*preQ1*) - for Q salvage. Q-modified tRNA is absent in $\Delta$*queD* and $\Delta$*queD* $\Delta$*yhhQ* strains grown in minimal media with glycerol; Q-modified tRNA is detected when a $\Delta$*queD* strain is grown in minimal media plus 10 nM *preQ0* or *preQ1* but is absent when a $\Delta$*queD* $\Delta$*yhhQ* strain is grown under these conditions. *yhhQ* expressed from a plasmid restores the presence of Q-modified tRNA in a $\Delta$*queD* $\Delta$*yhhQ* strain. | E,-, N |
| *yibJ* | 0.3 | No information about this protein was found by a literature search conducted on July 9, 2018. | E, N, N |
| *ymfK* | 0.4 | YmfK is a component of the relic lambdoid prophage e14 and is likely the SOS-sensitive repressor. It is similar to the P34 gene of the *Shigella flexneri* bacteriophage SfV and belongs to the LexA group of SOS-response transcriptional repressors. | E, E, E |

TABLE IV. **Below-threshold essential genes identified in *E. coli*.** This table describes the message numbers and annotations for essential genes that we estimated to have expression below the threshold of one message per cell cycle. However, in the final column, we show classifications from three different studies. Only one of the identified genes, *ymfK*, was consistently defined as essential.

However, since translation efficiency in yeast is not uniform, we must consider its variation for low-expression proteins. We estimate that the detection efficiency in yeast is roughly $10^3$ molecules. Using Eq. 15, we estimate that $\varepsilon \approx 100$ and our approximation holds at the low-expression detection limit.

In *E. coli*, the situation is somewhat more complicated. Unlike yeast, the translation efficiency is roughly constant (at high to intermediate expression levels) with respect to expression level [25], and therefore both terms in Eq. A57 are expected to scale like the canonical model ($\propto \mu_p^{-1}$). However, it is clear that the translation efficiency must significantly decrease for the lowest abundance proteins. This is visible even in Ref. [25] Fig. 1B, where the data falls below the predicted protein abundance at low message number. Note that these mass-spec measurements are not as sensitive as fluorescence-based measurements (*e.g.* only 64% proteome could be detected [58]). Furthermore, fits to the *E. coli* noise (Eq. A49) are consistent only with low values of $\varepsilon$. At sufficiently high expression levels such that we are confident about the translation efficiency, the noise is already very close to the noise floor.
